# Mother’s childhood adversity is associated with accelerated epigenetic aging in pregnancy and in male newborns

**DOI:** 10.1101/2023.03.02.530806

**Authors:** Christian K. Dye, Haotian Wu, Catherine Monk, Daniel W. Belsky, Daniel Alschuler, Seonjoo Lee, Kieran O’Donnell, Pamela Scorza

**Affiliations:** Department of Environmental Health Sciences, Columbia University, New York, New York, USA; Department of Psychiatry, Columbia University, Columbia University, New York, New York, USA; Division of Behavioral Medicine, New York State Psychiatric Institute, New York, New York, USA; Department of Epidemiology & Butler Columbia Aging Center, Columbia University, New York, New York, USA; Department of Biostatistics, Columbia University, New York, New York, USA; Yale Child Study Center, Yale School of Medicine, New Haven, Connecticut, USA; Department of Obstetrics and Gynecology, Columbia University, New York, New York, USA

**Keywords:** Epigenetics, epigenetic aging, adverse childhood experiences, adversity, DNA methylation, child development, maternal health, epigenetic clock, Horvath, Hannum, PhenoAge, GrimAge, DunedinPACE, aging, intergenerational

## Abstract

**Background:** Adverse childhood experiences (ACEs) are correlated with accelerated epigenetic aging, but it is not clear whether altered epigenetic aging from childhood adversities persists into adulthood and can be transmitted to the next generation. Thus, we tested whether mothers’ childhood adversity is associated with accelerated epigenetic aging during pregnancy and in their newborn offspring.

**Methods:** Data were from the Avon Longitudinal Study of Parents and Children (ALSPAC) sub-study, Accessible Resource for Integrated Epigenomic Studies (ARIES). Women provided retrospective self-reports during pregnancy of ACE exposure. DNA methylation was measured in mothers during pregnancy and cord blood at birth. Estimates of epigenetic age acceleration were calculated using Principal Components of Horvath, Hannum skin & blood, GrimAge, PhenoAge, and DunedinPACE epigenetic clocks for mothers; and the Knight and Bohlin cord blood clocks for newborns. Associations between a cumulative maternal ACE score and epigenetic age acceleration were estimated using linear regression models, adjusting for maternal age at pregnancy, smoking during pregnancy, education, and pre-pregnancy BMI. Models for offspring were stratified by sex and additionally adjusted for gestation age.

**Results:** Mothers’ total ACE score was positively associated with accelerated maternal PhenoAge and GrimAge. In newborn offspring, mothers’ total ACE score was positively associated with accelerated epigenetic aging in males using the Bohlin clock, but not in females using either epigenetic clock. We found male offsprings’ epigenetic age was accelerated in those born to mothers exposed to neglect using the Knight clock; and parental substance abuse using the Bohlin clock.

**Conclusion:** Our results show that mothers’ ACE exposure is associated with DNAm age acceleration in male offspring, supporting the notion that DNAm age could be a marker of intergenerational biological embedding of mothers’ childhood adversity. This is consistent with findings on vulnerability of male fetuses to environmental insults.

## 1. Introduction

Accumulating evidence suggests that negative early life experiences can have lasting effects on health and health behaviors throughout the life course. Adverse childhood experiences (ACEs) are traumatic experiences (*e*.*g*., abuse, neglect, and household dysfunction) that occur during childhood and adolescence and have been linked to premature morbidity and mortality.^1,2^ Notably, 45% of children in the US have experienced one or more types of ACEs and 10% have experienced at least three, making it a public health concern.^3^ Additionally, the impact of ACEs may affect multiple generations. A growing body of evidence has correlated maternal ACEs with offspring development, behavior, and health.^4-6^ It is believed that these intergenerational outcomes may be linked through other functional consequences of ACEs such as depression and anxiety. When these intermediate effects of ACEs occur during sensitive windows of development such as pregnancy, they may in turn influence fetal and early childhood development.^7^ Indeed, previous research has observed ACEs are associated with an increased risk of adverse mental health outcomes in mothers during pregnancy^8^ and psychosocial risk factors potentially mediate the effects of maternal ACEs on offspring development.^5,6^ However, although the relationship between parental, primarily maternal, ACEs and offspring health outcomes has been established, potentially through psychosocial and health risk behaviors, less is known regarding the underlying molecular mechanisms that may orchestrate the biological effects of this relationship.

DNA methylation, the addition of methyl groups on DNA nucleotides, is one such mechanism that may mediate the effects of maternal and fetal environmental exposures on offspring health.^9,10^ These modifications are responsive to environmental cues, are capable of altering cellular phenotypes (*i*.*e*., gene expression) without changes to the underlying genotype (*i*.*e*., genetics), and may underlie disease susceptibility.^11^ DNAm has garnered particular interest in biomedical research, given its application as a potential biomarker relating a broad spectrum of environmental influences to health and disease.^12^ Recently, DNAm patterns have been used as biomarkers to capture and approximate biological aging,^13^ which is the natural decline in tissue and cellular integrity, shifts in cellular activity, cellular senescence, and changes in normal physiological function over the lifespan.^14,15^ DNAm-based biomarkers of aging, or “epigenetic clocks”, have been shown to correlate with chronological age, and acceleration of epigenetic age has been shown to be associated with a spectrum of conditions, including unhealthy behaviors,^16^ frailty,^17^ cancer,^18^ diabetes,^19^ cardiovascular diseases,^20^ dementia,^21^ and mortality risk.^22^

Early life environmental conditions are known to alter the rate and direction of epigenetic aging.^23^ Evidence suggests that altered biological aging trajectories can be programmed early in development,^24^ even *in utero*, increasing disease risk over a person’s lifespan.^25^ Multiple studies have observed that ACEs are associated with accelerated epigenetic aging and adverse health outcomes in offspring later in life.^26-28^ Taken together, the existing evidence suggest that ACEs in mothers may accelerate the biological aging process in their offspring, especially as gestation represents a sensitive window of development for the offspring, which is consistent with evidence of premature morbidity and mortality in those directly exposed to ACEs.

In the current study, we sought to explore whether adverse experiences in mothers’ childhoods were associated with epigenetic age acceleration in mothers and in newborn infants. We leveraged DNAm data, collected from over 1,000 pregnant women and from umbilical cord blood from their babies at birth, in the Accessible Resource for Integrated Epigenetic Studies (ARIES),^29^ a sub-study of the Avon Longitudinal Study of Parents and Children (ALSPAC),^30^ in which 27% of participants reported experiencing at least one type of ACE, such as physical or emotional abuse or parental death, and 17% reported more than two.^31^ Leveraging rich DNAm data collected from mothers’ peripheral blood during pregnancy and newborns’ umbilical cord blood with ACE information collected from mothers, we examined the associations between maternal ACE exposure and epigenetic age acceleration in both the mothers and newborns.

## 2. Methods

### 2.1 Study Population

The Accessible Resource for Integrated Epigenomic Studies (ARIES) is a sub-study of the Avon Longitudinal Study of Parents and Children (ALSPAC), intended as a resource for investigating DNAm in health and child development.^29^ Briefly, ALSPAC recruited women who gave birth in the Avon Health District in the United Kingdom (UK) between April 1, 1991 and December 31, 1992, in which 13,761 pregnant women were eligible.^32^ The participation rate was between 85-90% of the eligible population and included families reflecting the general UK population, although less ethnic minority representation was unintentionally introduced (3% in ALSPAC vs. 7.6% in the UK). As a sub-study to ALSPAC, ARIES enrolled a subset of 1,018 mother-child dyads who had DNA samples collected from mothers at two time points (antenatal and follow-up at offspring mean age 15.5 years) and in children at three separate time points (birth, childhood [mean age 7.5 years], and adolescence [mean age 15.5 years]). DNA from children was collected from umbilical cord blood at birth and from peripheral blood at follow-up visits. DNA was collected from peripheral blood in mothers. Compared to the entire ALSPAC population, mothers in ARIES were older, more likely to have a non-manual profession, and less likely to have smoked while pregnant.^35^ Of the 1,018 mother-child pairs, we included mothers with non-missing ACE scores (pro-rated if < 3 individual ACE measurements were missing) and had antenatal and cord blood samples. In total, we retained 785 participants for maternal antenatal blood analyses and 753 newborns for umbilical cord blood analyses. Written informed consent was obtained from mothers for both themselves and participating children in ALSPAC and ARIES for study participation, including those used in this study. Ethical approval for this secondary data analysis was provided by the Columbia University Institutional Review Board.

### 2.2 Data Collection

Enrolled mothers submitted retrospective self-reports of their adverse childhood experiences (ACEs) between 18 and 32 weeks of gestation and when their child was approximately 3 years old.^33^ For consistency with the literature, we analyzed a cumulative measure of ten ACEs investigated in the landmark CDC-Kaiser ACE Study, the largest study of its kind that sought to identify a relationship between childhood ACEs and later-life health, and used often in ACE-related research.^1^ These ACE variables included emotional abuse, physical abuse, sexual abuse, emotional neglect, physical neglect, loss of a parent, domestic violence, family member with addiction, family member mental illness, and family member incarcerated. Rather than being included in ALSPAC as one “Adverse Childhood Experiences” questionnaire, relevant ACE questions were included separately in three different questionnaires administered to ALSPAC participants during pregnancy at approximately 18 and 32 weeks of gestation, or when their child was approximately three years old. Questionnaires were coded as binary variables (yes/no) for each ACE variable. A continuous variable summing the number of types of adversities reported (0-10) was created as a cumulative score of ACE exposure.

Maternal age was determined by the mother’s self-reported date of birth during the 8-week pregnancy interview. Parity (number of prior live or stillbirths) was ascertained by the mother’s self-report at 18 weeks gestation. Maternal smoking during pregnancy was reported by the mother at around 18 weeks gestation and calculated as the number of cigarettes smoked per day during the first trimester of pregnancy. Maternal education was determined by the mother’s self-report at around 32 weeks and classified as either less than high school, high school but no further education, some college or technical training, or completed university. Infant gestational age at birth was taken from obstetrics records. Pre-pregnancy BMI (kg/m^2^) was calculated using the mother’s self-reports of her pre-pregnancy weight and height from 12-week gestation. Approximately 44% of prenatal samples and 82% of cord blood samples were white blood cells (lymphocytes, monocytes, and granulocytes) while the remaining samples were either whole blood (approximately 56% of prenatal samples) or blood spots collected from umbilical cord blood (18% of cord blood samples). Because sample type was not correlated with the maternal ACE score (cord blood Wilcoxon test *p-value*= 0.35; antenatal blood *p*-value = 0.61), it was not used in downstream analyses.

### 2.3 DNA Methylation Preprocessing, Normalization, and Quantification

As part of the overall goal of ARIES, DNAm was assessed on peripheral blood samples collected from mothers during antenatal clinic visits and from umbilical cord blood samples taken at birth for newborns following standard protocols.^29^ DNA samples from maternal peripheral blood were collected at an average gestational age of 25.7±9.5 weeks. DNA was bisulfite-converted using the Zymo EZ DNA Methylation™ Kit (Zymo Research, Irvine, CA, USA). Illumina’s Infinium® HumanMethylation450 BeadChip (450K; Illumina, Inc., San Diego, CA, USA), covering >450,000 CpGs across 99% of RefSeq genes, was used for the accurate, reproducible, and reliable quantification of DNAm at cytosine-guanine dinucleotides (CpGs) across the genome at single-nucleotide resolution. Samples across all ARIES time points were semi-randomly distributed across 450K microarrays to minimize confounding by batch effects.

Initial quality control was performed using GenomeStudio (version 2011.1). Internal control probes on the 450K were used for initial quality control (QC) and recorded by ARIES staff. Raw IDAT files were preprocessed and normalized using the *meffil* package.^34^ First, samples whose average probe intensities were unreliable (detection *p*-value ≥ 0.01), thus failing initial QC, were repeated on the assay, and if still unsuccessful were excluded. Additionally, samples whose genotype information did not match with SNP data from the 450K annotation were excluded. For those samples that lacked genome-wide SNP data, we used sex prediction algorithms within the *meffil* package to accurately predict sex based on sex chromosome methylation patterns; those whose sex was mismatched between predicted sex based on DNAm data and actual known sex were removed as potentially mismatched samples. Further QC and preprocessing steps were performed, including failed CpG probes were removed (detection *p*-value ≥ 0.01 in 5% of participants or more) and sex chromosomes removed. Normalization was performed using the functional normalization method, a between-array normalization method extended from quantile normalization that removes technical variation in DNAm by using internal control probes to regress out variability explained by internal control probes of the 450K microarray.^35^ Chip effects were regressed on the raw betas before normalization and on the control matrix. Each sample was normalized individually to a cell type reference dataset using meffil.cell.count.estimates to estimate cell composition of samples.^40^ Finally, DNAm values (β-value) were generated for the remaining CpG loci, corresponding from a range between 0.0 (unmethylated) to 1.0 (completely methylated).

### 2.4 Epigenetic Clock Calculation

To calculate epigenetic age in blood samples from peripheral blood in mothers during pregnancy and cord blood in their newborn offspring, we used a modified Principal Component (PC) version of four of the most well-known epigenetic clocks, including the blood-based Hannum clock^36^ and pan-tissue Horvath clock,^37^ and two newer clocks that better predict lifespan and healthspan, GrimAge^38^ and PhenoAge.^39^ Due to the high variability across epigenetic clocks, we employed a PC-based approach to these epigenetic clocks recently described by Higgens-Chen *et al*. as a way to increase reliability of epigenetic clock results by removing technical noise, utilizing multiple sets of CpGs.^40^ This approach relies on utilizing CpG information to create PCs that are predictive of biological age. Notably, this method provides enhanced reliability for longitudinal data and better detection of associations with outcomes of interest. Epigenetic age of cord blood samples was calculated using the methods described in Bohlin *et al*.^41^ and Knight *et al*.^42^ In both maternal peripheral blood and newborn cord blood, epigenetic age acceleration was defined as the difference between DNAm age and chronological age. Age acceleration was defined as the resulting difference being positive, whereas a negative number signified epigenetic age deceleration. Similar to epigenetic age acceleration, we used the recently developed DunedinPACE,^43^ created using DNAm as a biomarker for assessing the pace of aging. Here, biological aging was considered accelerated if DunedinPACE > 1.0 and decelerated if < 1.0.

### 2.6 Statistical Analyses

Linear regression models were used with the sum score of maternal ACEs as the primary predictor and epigenetic age acceleration as the outcome. Analyses were conducted first with epigenetic age from maternal prenatal blood as the outcome and then repeated using epigenetic age from offspring umbilical cord blood as the predictor, testing each clock separately. Analyses for cord blood were stratified by infant sex, based on evidence that suggests sex differences in prenatal programming, including epigenetic alterations.^44, 45^ For all analyses, covariates included maternal age during pregnancy, parity, maternal smoking during pregnancy, maternal education, and pre-pregnancy BMI. For umbilical cord blood analyses, infant gestational age at birth was added as an additional covariate.

For the ACE items forming the 0-10 scale, we pro-rated missing items with the average of the non-missing items when < 3 items were missing. We treated individuals’ scores with > 2 ACE items missing as missing ACE scores. All covariates had less than 20% of missing values (0% - 4.1% missing-ness). We also tested each of the type of ACE separately as the primary predictor. In addition, we tested the following categories of ACEs: 1) Abuse (yes/no): presence of one or more of physical abuse, emotional abuse, sexual abuse; 2) Neglect (yes/no): presence of one or more of physical neglect and emotional neglect; and 3) Family adversity (yes/no): the presence of one or more of parental domestic violence, parental substance use, parental mental illness, parental death/separation.

We explored whether depression and anxiety were mediators between ACEs and biological aging, given previous evidence observing an association between ACEs and anxiety and depression;^44,45^ and depression and biological aging.^46^ In ALSPAC, the Edinburgh Postnatal Depression Scale (EPDS)^41^ was used to measure depression and anxiety symptoms in pregnant women at 18 and 32 weeks gestation. We used the summary score of the EPDS at 18 weeks gestation to test maternal depression/anxiety symptoms (EPDS score as a single variable) as a mediator in the association between maternal ACE score and epigenetic aging in maternal prenatal blood or offspring umbilical cord blood.

## 3. Results

### 3.1 Participant Characteristics

Descriptive statistics of study participants are provided in Table 1. The mean age (SD) of mothers (n=785) was 29.73 (4.3) years at 8 weeks of pregnancy. The average ACE score in the mothers was 1.27 (1.5), and 10.5% of women reported having experienced three or more ACEs. Among the mothers, depression and anxiety, as measured by the EPDS score, was on average 6.26 (4.6), and 17.97% of women reached an EPDS score cutoff (EPDS score > 10), which is recommended cutoff for a probable case of major depression in pregnant women.^47^ Parity, measured at 18 weeks gestation as the number of prior live or stillbirth, was 0.74 (0.8) for mothers. Of the 785 mothers, 686 (87.4%) did not smoke during pregnancy. Mothers who were smokers during pregnancy on average smoked 8.6 (7.1) cigarettes per day. Participants pre-pregnancy BMI was 22.7 (3.6) kg/m^2^. For education, most mothers had technical schooling or some college (37.50%), followed by high school (34.15%), university degrees (21.09%), and lastly, less than high school degree (7.25%). For newborns (n=753) of enrolled mothers, the average gestation age at birth was 39.59 (1.5) weeks.

**Table 1.** Descriptive Statistics of ALSPAC Study Participants

| Variables | Mean ± SD | Min./Max. |
| --- | --- | --- |
| ACE score | 1.27 ± 1.5 | 0/8 |
| EPDS score | 6.26 ± 4.6 | 0/25 |
| Maternal age (years) | 29.73 ± 4.3 | 16/42 |
| Parity | 0.74 ± 0.8 | 0/5 |
| Gestational age at birth (weeks) | 39.59 ± 1.5 | 30/44 |
| Pre-pregnancy BMI (kg/m<sup>2</sup>) | 22.7 ± 3.6 | 14.23/45.24 |
| Smoking during pregnancy (yes; %) | 12.4% | - |
| Number of cigarettes/day from smokers | 8.61 ± 7.1 | 1/30 |
| Maternal education (%) |  |  |
| Less than high school | 7.25 | - |
| High school | 34.15 | - |
| Technical/Some college | 37.5 | - |
| University | 21.09 | - |
| Baby sex (%) |  |  |
| Female | 51.39 | - |
| Male | 48.61 | - |

ALSPAC: Avon Longitudinal Study of Parents and Children; ACE: adverse childhood experiences; EPDS: Edinburgh Postnatal Depression Scale; BMI: body mass index.

### 3.2 Maternal ACEs and Epigenetic Aging in Mothers and Newborns

We estimated the association between the total maternal ACE score with epigenetic age acceleration using four different PC-based epigenetic clock measurements calculated from mothers’ peripheral blood taken at the mean gestation age (SD) of 25.7 (9.5) weeks of pregnancy and with two cord blood-based epigenetic clock measurements taken from newborns’ cord blood at birth at mean gestational age 39.59 (1.5) weeks. Here, we found maternal ACE score was associated with both maternal and offspring epigenetic age acceleration (Tables 2 and 3). A positive association was observed between mothers’ total ACE score and accelerated epigenetic aging based on PhenoAge (β [95% CI] =0.26 [0.01,0.51], *p*=.045) and GrimAge (β=0.25 [0.11,0.39], *p*<0.001) (Table 2). Although in the same direction, no significant associations were observed between total maternal ACE score and Horvath (β=0.15 [-0.07,0.37], *p*=0.18; Table 2) or Hannum (β=0.09 [-0.09,0.27], *p*=0.32), or DunedinPACE (β=0.00 [0.00,0.01], *p*=0.24).

**Table 2.** Association Between Cumulative ACEs and Epigenetic Age Acceleration in Pregnancy

| DNAm Clock | β-Coefficient | 95% CI | *p*-value |
| --- | --- | --- | --- |
| Horvath | 0.15 | (-0.07,0.37) | 0.18 |
| Hannum | 0.09 | (-0.09,0.27) | 0.32 |
| PhenoAge | 0.26 | (0.01, 0.51) | 0.05 |
| GrimAge | 0.25 | (0.11,0.39) | <0.001 |
| DunedinPACE | 0 | (0.00,0.01) | 0.24 |

All models were adjusted for maternal age (during pregnancy), parity, smoking status (during pregnancy), education, and pre-pregnancy BMI.

**Table 3.** Association Between Maternal Cumulative ACEs and Epigenetic Age Acceleration in Newborns

| DNAm Clock | Male |  |  | Female |  |  |
| --- | --- | --- | --- | --- | --- | --- |
| | β-Coefficient | 95% CI | *p*-value | β-Coefficient | 95% CI | *p*-value |
| Knight | 0.12 | (-0.03,0.26) | 0.11 | -0.07 | (-0.19,0.04) | 0.22 |
| Bohlin | 0.06 | (0.00, 0.11) | 0.04 | 0.01 | (-0.04,0.07) | 0.57 |

All models were adjusted for maternal age (during pregnancy), parity, smoking status (during pregnancy), education, and pre-pregnancy BMI.

In offspring umbilical cord blood, mothers’ total ACE score was associated with accelerated epigenetic aging using the Bohlin epigenetic clock in males (β=0.06 [0.001,0.11], *p*= 0.04; Table 3), but not females (β=0.01 [-0.04,0.07], *p*=0.57). No associations were found between maternal total ACEs and the Knight epigenetic clock in either sex.

### 3.3 Depression and Anxiety as Mediator between ACEs and Epigenetic Aging in Mothers and Newborns

We sought to determine whether depression and anxiety mediates the effects of maternal ACEs on epigenetic age acceleration in both mothers and their newborns. However, maternal depression and anxiety symptoms, measured using the EPDS score, were not associated with DNAm age acceleration in either maternal peripheral blood during pregnancy or cord blood from newborn males (*p*-value range: 0.22 - 0.84). We therefore did not perform mediation analyses.

### 3.4 Individual ACEs and Epigenetic Aging in Mothers and Newborns

In addition to the cumulative score of ACEs from mothers, we sought to identify which individual ACEs may be associated with epigenetic age acceleration in both mothers’ peripheral blood during pregnancy and newborn offspring cord blood. Here, we found multiple types of ACEs were positively associated with accelerated epigenetic aging using the GrimAge clock in maternal peripheral blood (Figure 1. A-C. Supplementary Table 1), including emotional abuse (β=0.82 [0.05,1.60], *p*=0.04), physical neglect (β=2.43 [0.80,4.06], *p*=0.003), parental substance abuse (β=0.98 [0.20,1.77], *p*=0.01), family mental illness (β=0.49 [0.00,0.98], *p*=0.05), and in the categories of any abuse (β=0.68 [0.23,1.13], *p*=0.003) and any type of family adversity (β=0.51 [0.08,0.94], *p*=0.02). Likewise, we observed an association between accelerated epigenetic aging using the PhenoAge clock in maternal peripheral blood (Figure 1. A-C. Supplementary Table 1) with 2 individual ACEs: physical abuse (β=2.30 [0.21,4.39], *p*=0.03), and family mental illness (β=0.89 [0.01,1.78], *p*=0.05) and the any abuse ACE category (β=0.90 [0.08,1.71], *p*=0.03). Using the pace of aging DNAm clock, DunedinPACE, we observed a deceleration in biological aging that was associated with physical abuse in mothers (β=0.05 [0.00,0.10]; *p*=0.04). In male newborns’ cord blood analyses, we found specific maternal ACEs were associated with epigenetic age acceleration using the Knight clock (Figure 2. Supplementary Table 2), including emotional neglect (β=0.62 [0.09,1.15], *p*=0.02) and any type of neglect (β=0.58 [0.05,1.11], *p*=0.03). Using the Bohlin clock, we found parental substance abuse was associated with epigenetic age acceleration in newborn males (β=0.34 [0.03,0.65], *p*=0.03; Figure 2. Supplementary Table 2). In female newborns’ cord blood, we observed parental domestic violence was associated with decelerated epigenetic aging using the Knight clock (β=-0.67 [-1.25,-0.08], *p*=0.03).

**Figure 1.**
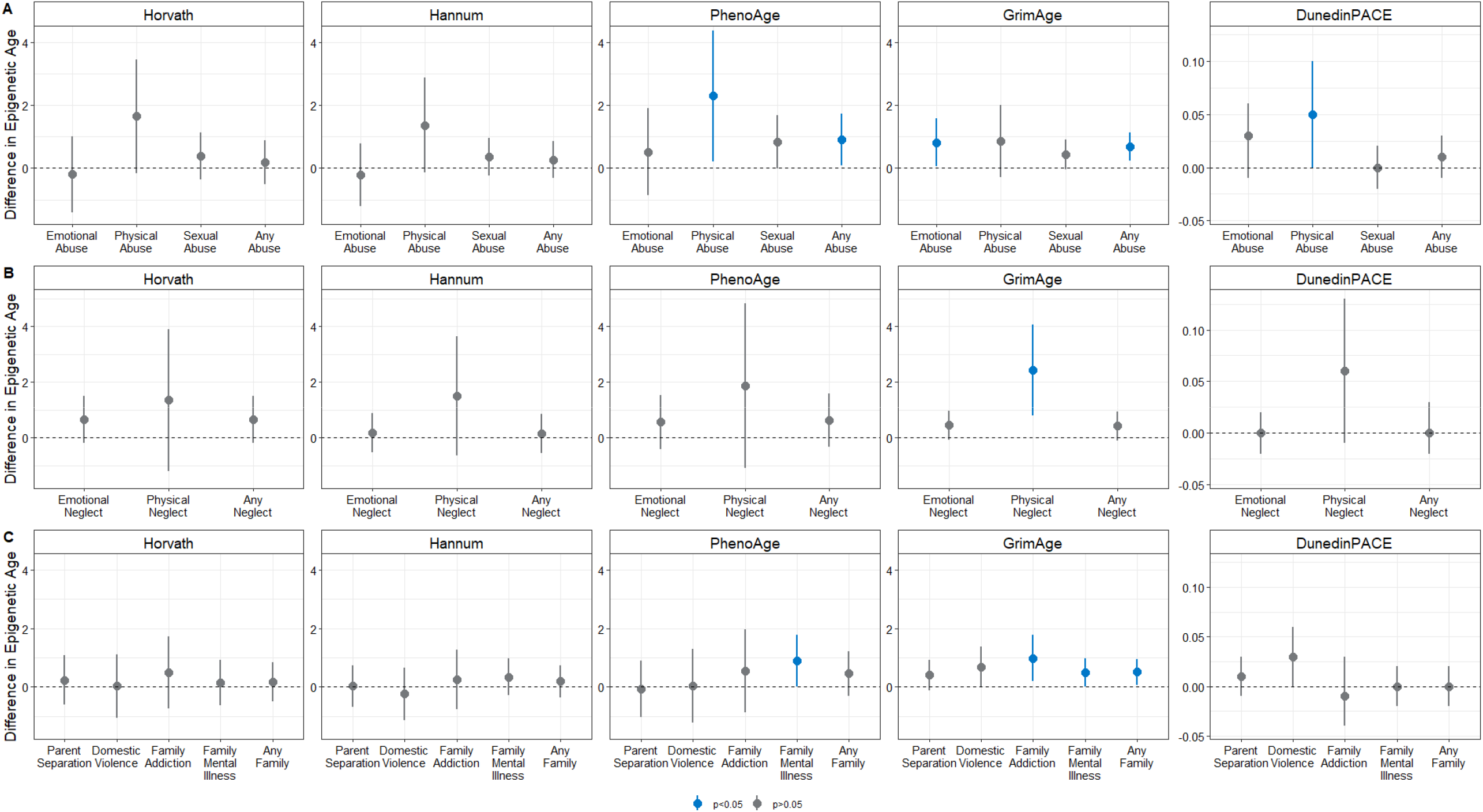
Individual maternal ACEs and change in epigenetic age deviations in pregnant women. Effect estimates for the change in epigenetic age in peripheral blood from pregnant women, calculated using 5 DNAm clocks: Horvath, Hannum, PhenoAge, GrimAge, and DunedinPACE. **A**. Relative change in epigenetic age for exposure to abuse (left to right, each plot): emotional abuse, physical abuse, sexual abuse, or any type of abuse as a child. **B**. Relative change in epigenetic age for exposure to neglect (left to right, each plot): emotional neglect, physical neglect, any neglect. **C**. Relative change in epigenetic age for exposure to household dysfunction (left to right, each plot): parent separation, domestic violence, family addiction (substance abuse), family mental illness, any family dysfunction. Blue color schemes represent statistical significance (*p*<0.05).

**Figure 2.**
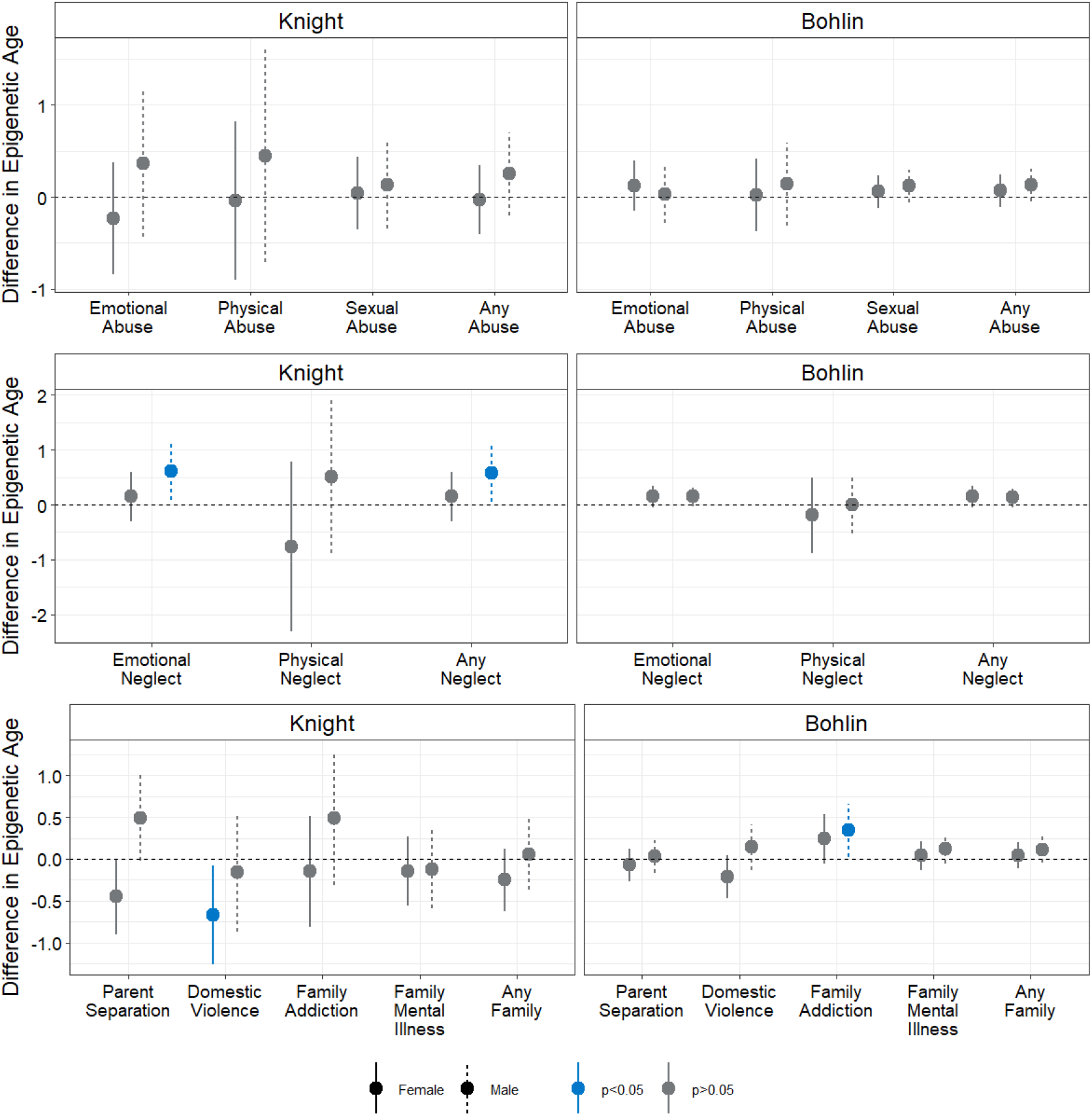
Individual maternal ACEs and change in epigenetic age deviations in newborn offspring. Effect estimates for the change in epigenetic age in cord blood collected from newborns, calculated using two clocks: Knight (left figures) and Bohlin (right figures). **A**. Relative change in epigenetic age for exposure to abuse (left to right, each plot): emotional abuse, physical abuse, sexual abuse, or any type of abuse as a child. **B**. Relative change in epigenetic age for exposure to neglect (left to right, each plot): emotional neglect, physical neglect, any neglect. **C**. Relative change in epigenetic age for exposure to household dysfunction (left to right, each plot): parent separation, domestic violence, family addiction (substance abuse), family mental illness, any family dysfunction. Female newborns represented by solid lines; male newborns represented by dashed line. Blue color schemes represent statistical significance (*p*<0.05).

## 4. Discussion

The ALSPAC study provides a unique opportunity to investigate the intra- and intergenerational effects of maternal ACEs, while the ARIES sub-study affords the opportunity to gain insight into the potential molecular mechanisms, including epigenetic modifications that underlie the observed relationship. In this study we sought to identify whether ACEs were associated with differences in epigenetic age in women during pregnancy and in their newborn offspring. Our results show that mothers’ cumulative ACE exposure was significantly and positively associated with PhenoAge and GrimAge acceleration in maternal prenatal peripheral blood and the Knight clock in newborn male cord blood, but not newborn females. These results suggest the biological embedding of ACEs via epigenetic aging and supports the hypothesis that intergenerational transmission of adversity can manifest through early-life changes in epigenetic aging. This is notable, as epigenetic aging is a potential indicator of negative health and developmental trajectories for offspring.^48^ Epigenetic aging in children has been associated with alterations in brain volumes, cortical thickness, cortical surface areas and abnormalities in neuronal microstructure in a range of brain regions; as well as a range of other offspring developmental and behavioral outcomes.^49^

In data from ALSPAC, cumulative and individual ACE exposure of the offspring themselves during childhood (aged 0-14 years), including emotional and physical abuse, were linked to epigenetic age acceleration through 17 years of age, with stronger effect estimates in females.^50^ Our study, looking at the effect of ACE exposure in the previous generation (i.e. the mothers) provided generally consistent results, as physical and emotional abuse in mothers’ own childhoods were associated with PhenoAge and GrimAge acceleration, respectively, during pregnancy. However, we also observed that other forms of ACEs, such as physical neglect, family mental illness, and family addiction, were also related to PhenoAge and GrimAge in the mothers.

In terms of the different epigenetic clocks, we observed significant relationships between the number and type of ACEs with epigenetic age acceleration in women during pregnancy, measured using PCs of both the PhenoAge and GrimAge DNAm clocks, but not with Horvath or Hannum DNAm clocks. Consistent with these findings, previous evidence found cumulative and individual ACEs were associated with accelerated epigenetic aging, measured by GrimAge^51^ and PhenoAge.^27^ The lack of association between maternal ACEs and epigenetic aging using the first-generation clocks (*e*.*g*., Hannum and Horvath clocks) may arise from the limitations of these clocks in predicting disease phenotypes and all-cause mortality, as these clocks were intended, and trained, as predictors of chronological age.^36,52^ The second-generation clocks (*e*.*g*., PhenoAge and GrimAge) were developed utilizing DNAm data to identify relationships with clinical biomarkers of aging, age-related disease risk, and mortality.^38,39^ Thus, the significant associations observed between cumulative and individual ACEs with PhenoAge and GrimAge, but not Horvath and Hannum clocks, may be due to the training of these clocks to perform better as predictors of clinical phenotypes of aging, age-related disease risk, and mortality.^39,53^ Notably, we did not observe evidence that emotional and physical abuse led to differences in epigenetic aging of the newborns, although it is possible that the intergenerational aging influence of those ACEs are not observable until later in life. Our findings did show that other ACEs, such as neglect and parental substance abuse, were associated with accelerated epigenetic aging in in male newborns but not females. This is consistent with other studies that reported maternal ACEs prior to pregnancy were associated with longer telomere length and accelerated epigenetic aging in children several years after birth (at ages, 7, 9, and 14), although whether this was linked to gestational age acceleration was not investigated.^54^

Mechanistically, early life adversities are likely embedded early in life and contribute to long-term changes in health behaviors (*e*.*g*., diet, smoking, physical activity) and psychosocial responses (*i*.*e*., stress response, depression, anxiety).^55,56^ Eventually, they can have tremendous downstream physiological effects on the affected individual, such as changes in biological functions like the hypothalamic-pituitary-adrenal (HPA) axis and sympathetic nervous system, chronic inflammatory phenotypes, epigenetic alterations, and neural effects, which consequently can lead to pathophysiological states underlying age-related disease susceptibility.^57-59^ When this cascade of events from early life adversity to biological changes occurs during sensitive windows of development, such as pregnancy, they may impart similar effects on the developing fetus.^5^ Notably, depression and anxiety symptoms did not mediate the relationship between maternal ACEs and epigenetic aging in our study. This is in contrast to previous studies that have found depression in pregnancy to be associated with DNAm age deviations in the infant at birth in umbilical cord blood^42,60-62^ and maternal prenatal anxiety to be predictive of childhood epigenetic age acceleration.^63^ Interestingly, the effects in the latter study were largely observed in males but not females, supporting our findings of sex differences in maternal ACE-associated age acceleration.

The Developmental Origins of Health and Disease (DOHaD) hypothesis suggests that exposure to environmental factors (*e*.*g*., environmental toxicants, malnutrition, psychosocial risk behaviors, *etc*.) during labile windows of development, such as antenatal and postnatal periods, may contribute to offspring disease susceptibility later in life^64^, including cardiovascular diseases, metabolic diseases, cancer, and autoimmune diseases.^65-68^ Indeed, due to the biological connection between mother and offspring, including the placenta, maternal exposures may be either directly transmitted to or have indirect biological effects that are conferred to the developing neonate.^69,70^ Interestingly, maternal ACEs have been associated with adverse environmental exposures during pregnancy (*i*.*e*., depressive symptoms, stress, anxiety) that precede adverse health and developmental outcomes in offspring, supporting the DOHaD hypothesis in the intergenerational transmission of disease susceptibility.^71,72^ Epigenetic mechanisms provide a fundamental molecular mechanism that supports the DoHAD hypothesis, given epigenetic alterations are environmentally responsive, regulate cellular phenotypes, are amenable to reprogramming during fetal development in response to altered fetal environments, and may be a biomarker of age-related disease risk.^73^ Epigenetic alterations have been observed at stress-related genes from neonates of mothers exposed to ACEs.^74^ Perinatal exposure to adverse environmental conditions may accelerate epigenetic aging at birth^75^, and exposomic analyses during prenatal and early childhood observed accelerated epigenetic aging in children.^76^ Given the role for epigenetic mechanisms in facilitating the effects of our environment on cellular phenotype, including fetal development trajectories, it is likely that the effects of early life adversities can have their most pronounced effects later in life particularly during sensitive windows of development - where the DoHAD hypothesis is most relevant (*i*.*e*., pregnancy and early life).

While this is the first study, to our knowledge, to report the association between mothers’ ACE exposure and epigenetic aging in both mothers during pregnancy and their newborn offspring, results should be interpreted in the context of several limitations that should be noted. The prevalence of ACEs in the ALSPAC sample is relatively low, though this is the case for much existing literature on the topic. Studies in samples exposed to relatively low levels of adversity might fail to detect associations, particularly those with small effect sizes as is the case with psychosocial exposures and intergeneration outcomes. Additionally, because our analysis was not performed in a population highly exposed to adversities, our findings might not generalize to such populations. Only 3% of ALSPAC mothers self-identified as belonging to a racial/ethnic minority group. Unfortunately, marginalized populations are those most affected by intergenerational transmission of sequela of childhood adversity. Marginalized racial and ethnic groups, including Black and Latinx populations, sustain a higher exposure to and experience more ACE’s than their White counterparts.^77^ Future research should therefore focus on populations with very high exposure to adversity.

Taken together, epigenetic age acceleration observed in newborn offspring, specifically males, may be the result of the early life adversities of exposed mothers that had intragenerational biological (*e*.*g*., psychosocial and health behaviors, chronic stress, *etc*.) and molecular effects (*i*.*e*., biological age acceleration), and if they persisted through pregnancy were potentially embedded in newborn offspring.

## 5. Conclusions

In this study, we found that the accumulation of adverse childhood experiences in women were associated with accelerated biological aging – as compared to chronological age – later in life. This appears to be driven by a variety of types of ACEs, including physical abuse, emotional abuse, physical neglect, family mental illness, and family addiction. Further, our results suggests that maternal adversity early in life is associated with acceleration in biological aging in male, but not female, newborns at birth. To our knowledge, this is the first study to report differences in biological aging in both pregnant women who have been exposed to early life childhood adversities as well as in their newborn offspring. Future studies should confirm these findings elsewhere and in a more diverse cohort, including underrepresented minorities and others most at risk for adverse childhood experiences, elucidate the long-term health effects of intergenerationally transmitted biological aging in the offspring, and further explore the mediating mechanisms between maternal adversity during childhood and changes in biological aging.

## Authorship Contribution Statement

**Christian K. Dye:** manuscript authorship (writing, reviewing, editing, figures/tables/legends). **Haotian Wu:** manuscript authorship (writing, reviewing, figures, and editing). **Catherine Monk:** intellectual resource for formal analyses and framework for study. **Daniel W. Belsky:** intellectual resource for DNAm analysis and framework. **Daniel Alschuler:** formal analyses, data access, manuscript authorship (reviewing and editing). **Seonjoo Lee:** intellectual resource for formal analyses. **Kieren O’Donnell:** intellectual resource for formal analyses. **Pamela Scorza:** conceptualization, manuscript authorship (writing, reviewing, editing), supervision.

## Acknowledgements

The authors would like to thank the Avon Longitudinal Study of Parents and Children participants and staff, without your time, dedication, and help, none of this work would have been made possible. This study was supported by the National Institute of Environmental health Sciences grant: T32ES007322 (CKD) and the National Institute of Mental Health grant K01MH117443 (PS). The Avon Longitudinal Study of Parents and Children is funded by the Medical Research Council, the Wellcome Trust and the University of Bristol. The Accessible Resource for Integrated Epigenomics Studies is funded by the Biotechnology and Biological Sciences Research Council.

---

**Supplementary Table 2.**
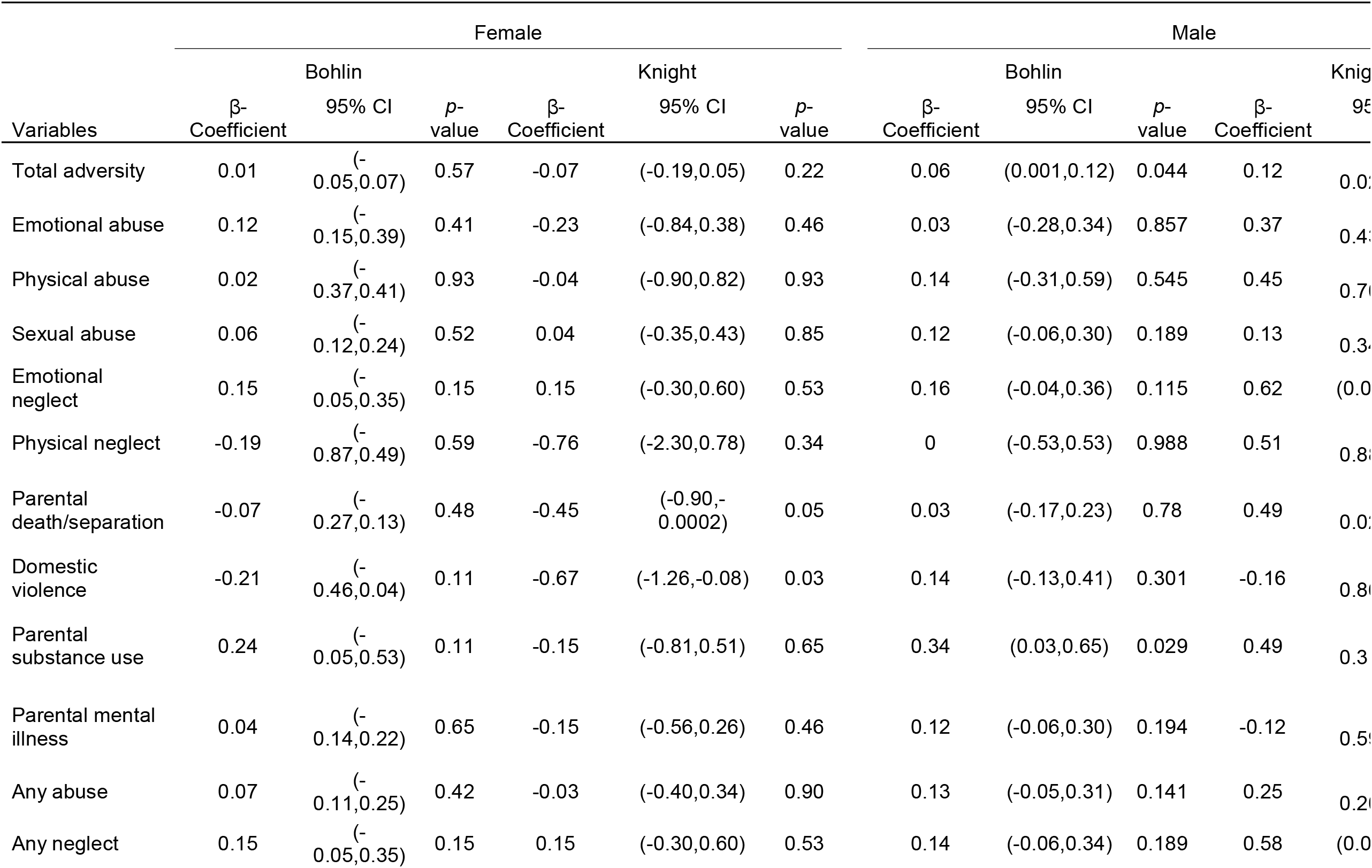
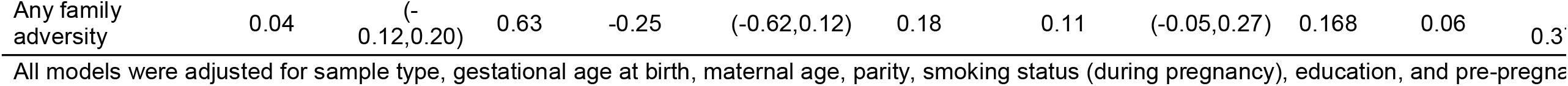
Individual maternal ACEs and epigenetic age acceleration in newborns.

