## Supplementary Table 1 for "Mother’s childhood adversity is associated with accelerated epigenetic aging in pregnancy and in male newborns"

**Supplementary Tables**

| **Supplementary Table 1. Individual maternal ACEs and epigenetic age acceleration in pregnant mothers.** | | | | | | |
| --- | --- | --- | --- | --- | --- | --- |
|  | PhenoAge | | | GrimAge | | |
| Variables | β-Coefficient | 95% CI | *p*-value | β-Coefficient | 95%CI | *p*-value |
| Total adversity | 0.26 | (0.01,0.51) | 0.05 | 0.25 | (0.11,0.39) | <0.001 |
| Emotional abuse | 0.52 | (-0.88,1.92) | 0.47 | 0.82 | (0.05,1.60) | 0.04 |
| Physical abuse | 2.3 | (0.21,4.39) | 0.03 | 0.86 | (-0.29,2.02) | 0.14 |
| Sexual abuse | 0.83 | (-0.02,1.68) | 0.06 | 0.43 | (-0.04,0.91) | 0.07 |
| Emotional neglect | 0.56 | (-0.41,1.53) | 0.25 | 0.45 | (-0.09,0.98) | 0.1 |
| Physical neglect | 1.87 | (-1.09,4.83) | 0.22 | 2.43 | (0.80,4.06) | 0.003 |
| Parental death/separation | -0.08 | (-1.05,0.89) | 0.88 | 0.4 | (-0.14,0.93) | 0.15 |
| Domestic violence | 0.04 | (-1.22,1.30) | 0.96 | 0.68 | (-0.01,1.37) | 0.06 |
| Parental substance use | 0.54 | (-0.88,1.96) | 0.45 | 0.98 | (0.20,1.77) | 0.01 |
| Parental mental illness | 0.89 | (0.01,1.78) | 0.05 | 0.49 | (0.00,0.98) | 0.05 |
| Any abuse | 0.9 | (0.08,1.71) | 0.03 | 0.68 | (0.23,1.13) | 0.003 |
| Any neglect | 0.62 | (-0.34,1.59) | 0.21 | 0.42 | (-0.11,0.96) | 0.12 |
| Any family adversity | 0.46 | (-0.32,1.23) | 0.25 | 0.51 | (0.08,0.94) | 0.02 |
| All models were adjusted for maternal age, parity, smoking status (during pregnancy), education, and pre-pregnancy BMI. | | | | | | |
