## Supplementary Table 2 for "Mother’s childhood adversity is associated with accelerated epigenetic aging in pregnancy and in male newborns"

| **Supplementary Table 2. Individual maternal ACEs and epigenetic age acceleration in newborns.** | | | | | | | | | | | | |
| --- | --- | --- | --- | --- | --- | --- | --- | --- | --- | --- | --- | --- |
|  | Female | | | | | | Male | | | | | |
|  | Bohlin | | | Knight | | | Bohlin | | | Knight | | |
| Variables | β-Coefficient | 95% CI | *p-value* | β-Coefficient | 95% CI | *p-value* | β-Coefficient | 95% CI | *p-value* | β-Coefficient | 95% CI | *p-value* |
| Total adversity | 0.01 | (-0.04,0.07) | 0.57 | -0.07 | (-0.19,0.04) | 0.22 | 0.06 | (0.00,0.11) | 0.04 | 0.12 | (-0.03,0.26) | 0.11 |
| Emotional abuse | 0.12 | (-0.16,0.39) | 0.41 | -0.23 | (-0.85,0.38) | 0.46 | 0.03 | (-0.28,0.34) | 0.86 | 0.37 | (-0.44,1.18) | 0.37 |
| Physical abuse | 0.02 | (-0.37,0.41) | 0.93 | -0.04 | (-0.91,0.83) | 0.93 | 0.14 | (-0.31,0.58) | 0.55 | 0.45 | (-0.70,1.61) | 0.44 |
| Sexual abuse | 0.06 | (-0.12,0.24) | 0.52 | 0.04 | (-0.36,0.43) | 0.85 | 0.12 | (-0.06,0.30) | 0.19 | 0.13 | (-0.34,0.60) | 0.59 |
| Emotional neglect | 0.15 | (-0.05,0.36) | 0.15 | 0.15 | (-0.31,0.61) | 0.53 | 0.16 | (-0.04,0.37) | 0.12 | 0.62 | (0.09,1.15) | 0.02 |
| Physical neglect | -0.19 | (-0.89,0.50) | 0.59 | -0.76 | (-2.31,0.79) | 0.34 | 0 | (-0.53,0.54) | 0.99 | 0.51 | (-0.89,1.91) | 0.47 |
| Parental death/separation | -0.07 | (-0.28,0.13) | 0.48 | -0.45 | (-0.91,0.00) | 0.05 | 0.03 | (-0.17,0.23) | 0.78 | 0.49 | (-0.02,1.00) | 0.06 |
| Domestic violence | -0.21 | (-0.47,0.05) | 0.11 | -0.67 | (-1.25,-0.08) | 0.03 | 0.14 | (-0.13,0.42) | 0.3 | -0.16 | (-0.87,0.55) | 0.65 |
| Parental substance use | 0.24 | (-0.06,0.54) | 0.11 | -0.15 | (-0.83,0.52) | 0.65 | 0.34 | (0.03,0.65) | 0.03 | 0.49 | (-0.32,1.30) | 0.24 |
| Parental mental illness | 0.04 | (-0.14,0.23) | 0.65 | -0.15 | (-0.57,0.26) | 0.46 | 0.12 | (-0.06,0.30) | 0.19 | -0.12 | (-0.59,0.36) | 0.63 |
| Any abuse | 0.07 | (-0.10,0.24) | 0.42 | -0.03 | (-0.40,0.35) | 0.9 | 0.13 | (-0.04,0.30) | 0.14 | 0.25 | (-0.19,0.70) | 0.26 |
| Any neglect | 0.15 | (-0.05,0.36) | 0.15 | 0.15 | (-0.31,0.61) | 0.53 | 0.14 | (-0.07,0.34) | 0.19 | 0.58 | (0.05,1.11) | 0.03 |
| Any family adversity | 0.04 | (-0.12,0.21) | 0.63 | -0.25 | (-0.62,0.12) | 0.18 | 0.11 | (-0.05,0.28) | 0.17 | 0.06 | (-0.37,0.48) | 0.79 |
| All models were adjusted for gestational age at birth, maternal age, parity, smoking status (during pregnancy), education, and pre-pregnancy BMI. | | | | | | | | | | | | |
